# Robust enhancement of motor sequence learning with 4mA transcranial electric stimulation

**DOI:** 10.1101/2022.06.07.495056

**Authors:** Gavin Hsu, A. Duke Shereen, Leonardo G. Cohen, Lucas C. Parra

## Abstract

**Background and Objectives:** Motor learning experiments with transcranial direct current stimulation (tDCS) at 2mA have produced mixed results. We hypothesize that tDCS will boost motor learning provided sufficiently high field intensity on the motor cortex.

**Methods:** In a single-blinded, between-subject design, 72 healthy right-handed participants received either anodal or cathodal tDCS at 4mA while they learned to perform a sequence of key presses using their non-dominant hand for about 12 minutes. Cathodal stimulation served as an active control for sensation. A separate sham-stimulation group established baseline performance. Gains during practice and rest periods were analyzed (called micro-online and -offline learning). The target for stimulation was identified on the motor cortex using fMRI. After optimization with individual current flow models, we selected a single montage for all 108 participants with 4 frontal and 4 parietal electrodes each drawing 1mA.

**Results:** We found significant gains in performance with anodal stimulation (Cohen’s d=0.7). The boost in performance persisted for at least one hour. Subsequent learning for a new sequence and the opposite hand also improved. Concurrent tDCS enhanced micro-offline learning, while subsequent learning relied on micro-online gains. Sensation ratings were comparable in the active groups and did not exceed moderate levels. The new electrode montage achieved a better tradeoff between stimulation intensity and sensations on the scalp as compared to alternative montages.

**Conclusion:** The present paradigm shows reliable behavioral effects at 4mA and is well-tolerated. It may serve as a go-to experiment for future studies on motor learning and tDCS.

**Highlights:**

- tDCS resulted in a lasting boost of concurrent learning with effect size of Cohen’s d=0.7.
- Subsequent learning was also improved, indicating a form of meta-learning.
- Detailed analysis of behavior suggests an effect of tDCS on sequence consolidation.
- A novel electrode montage with 1mA through each of 4+4 electrodes was well-tolerated.

## Introduction

Transcranial direct current stimulation (tDCS) has been shown to affect neuronal excitability^1,2^, and some have argued that this effect may result from a modulation of synaptic efficacy^3^. We have determined *in vitro* that direct current stimulation (DCS) can interact with endogenous plasticity mechanisms to boost synaptic enhancements^4–6^. The implication is that tDCS may boost learning in tasks where learning is known to result from changes in synaptic efficacy, including motor skill learning tasks in both humans and rodents ^7–9^. Indeed, a number of human behavioral studies paired tDCS with a motor learning protocol with the aim of enhancing learning effects. However, they have produced mixed results^10^. Several studies reported that concurrent anodal stimulation over the primary motor cortex (M1) or the cerebellum enhances explicit motor sequence learning ^11–14^. Others found that cerebellar tDCS at 2mA did not affect learning performance, and did not change neuronal activity recorded with functional magnetic resonance imaging (fMRI)^15,16^. Yet others reported a monotonic increase of motor learning performance when increasing stimulation intensity to 4mA ^17^, but failed to control for sensation that increases with intensity^18^. In total, we are still lacking a dependable behavioral method for assessing efficacy of tDCS in modulating motor learning.

This lack of reliability may be due in part to an inadequate strength of stimulation. Electric fields induced on the cortical surface with conventional tDCS are less than 1V/m^19^. This is well below values used *in vitro* to demonstrate effects on plasticity ^4,5,20–23^. The objective of this study is to establish a go-to behavioral experiment to test the effects of tDCS on learning. We hypothesize that motor sequence learning can be reliably enhanced with concurrent tDCS by increasing the current intensity to 4mA from the conventional 2mA (H1). Furthermore, we expect this boosting effect to last over time (H2). Informed by our prior *in vitro* DCS work, we hypothesize that the lasting tDCS effect is specific to the hemisphere paired with stimulation (H3) and specific to the sequence paired with stimulation (H4).

We test this using a classic motor sequence learning task, namely the “finger tapping task” (FTT). Performance gains in this task were shown to correlate with functional changes in M1^24^. When combined with concurrently applied anodal stimulation, we predict a boost in task performance. The stimulation specifically targets the “hand knob”, which we localize with fMRI. We optimized placement of high-definition Tdcs (HD-tDCS) electrodes to maximize intensity on this target while distributing currents through 4 electrode pairs each passing 1mA. Consequently, the benefits of this setup are twofold: there is better spatial targeting^25^ as well as a reduction in discomfort from spreading out currents on the scalp^26–28^. To control for sensation effects, which are inevitable at 4mA, we compare anodal vs. cathodal stimulation in two different cohorts, as well as a no-stimulation control group. We find a robust increase in performance with anodal stimulation lasting up to 1 hour after stimulation, as well as gains in learning new sequences after stimulation.

## Methods

### Summary

We aimed to measure the modulatory effects of tDCS behaviorally through the FTT. Firstly, we determined optimal electrode placement to maximize the electric field induced at the brain region most active during the learning task. Through fMRI with a small sample of 10 subjects we located stimulation targets on the “hand knob” region of M1, which were entered as targets in current flow models of the individual brains to generate a common electrode montage that would be used for all subjects (N=108) during the motor sequence learning task. The montage would consist of 4 electrode pairs, with 4 anodes plus 4 cathodes (4+4). For the learning task subjects were divided into three groups (N=36 each), receiving anodal stimulation, active sham with cathodal stimulation, and no stimulation. Here, “anodal” stimulation is defined as inward current on the targeted cortical structure, and polarity is reversed in “cathodal” stimulation.

Stimulation was applied during FTT performance (H1), followed after an hour-long break by repetitions of the FTT to determine carryover effects across time (H2), brain hemispheres (H3), and different sequences (H4).

### Target Location

Target location and electrode placement were determined on an imaging cohort of 10 healthy adults (2 female, 8 male, age range = 18–55 years, mean ± SD = 24.2 ± 6.12 years). All participants provided written consent to participate in this research, under approval of the City University of New York Institutional Review Board (IRB). Exclusion criteria for potential participants included any history of neurological or psychiatric disorders, traumatic brain injury, disabilities in the upper extremities, severe visual impairment, and any MRI contraindication.

Participants were scanned in a Siemens 3T Prisma MRI (Siemens, Munich, Germany) at the City University of New York’s Advanced Science Research Center using a 32 channel head coil receiver. The data for each participant consisted of a sagittal three-dimensional T1-weighted magnetization-prepared rapid gradient echo (MPRAGE) anatomical scan and a functional echo planar imaging scan during a hand-motor task. The anatomical scans were collected with the following parameters: 208 slices, TR/TE/TI = 2400/2.15/1000 ms, slice thickness = 1 mm, flip angle = 8 degrees, FOV = 256×240, in-plane resolution = 256×240 mm^2, and acquisition time = 5:42 min. Task fMRI were collected using an echo-planar imaging scan: 60 interleaved axial slices, slice thickness = 2.4 mm, no slice gap, multiband factor = 6, TR/TE = 800/30 ms, flip angle = 52 degrees, FOV = 216×216 mm^2, in-plane resolution = 90 × 90, and acquisition time = 6:54 min. During task fMRI, participants held one 4-button response pad in each hand, with one finger on each button. Prompted by a computer monitor, the participants pressed buttons in alternating sets of 30 seconds on one hand at a time, alternating between hands with 30 seconds of rest between sets. To simulate the FTT, during each set participants pressed the buttons one at a time in sequence from the pinky toward the index finger and repeated this sequence as many times as possible during the 30 seconds. 10 sets were completed for each hand, for approximately 10 minutes total during the fMRI session. Contrasts in blood oxygenation level dependent (BOLD) signal intensity during performance of finger tapping were calculated in AFNI (Analysis of Functional NeuroImages) relative to the signals during the rest periods (separately for left and right hands). Subject level preprocessing included: slice time correction, despiking, coregistering the functional time series to a target volume and aligning the anatomical to the coregistered data, censoring data with motion greater than 1 mm from the target volume, smoothing the fMRI by applying a 5 mm Gaussian kernel to the brain extracted data, bandpass filtering between 0.01 and 0.1 Hz, and detrending the time series to correct for signal drift. To remove artifact from signal, time series data from the white matter and cerebral spinal fluid, along with 6 motion regressors from the registration were included as nuisance regressors in AFNI’s ‘3dDeconvolve’ function which modeled the data to the input stimulus (left/right/rest finger tapping blocks) and produced statistical maps of the regression beta coefficients and t-statistics for significance of the coefficients. The resulting t-statistic maps were thresholded to determine the pixels most active during sequential finger movement (Fig. 1, for left hand). We focus on the left hand (right hemisphere) as the sequence learning task will target the non-dominant hand in right-handed participants. Cortical parcellation was performed using FreeSurfer^29,30^, which we used to narrow down active regions to the “hand knob” area on the precentral gyrus, and within those boundaries manually selected a single voxel with maximal or near-maximal statistical value as the target.

**Figure 1.**
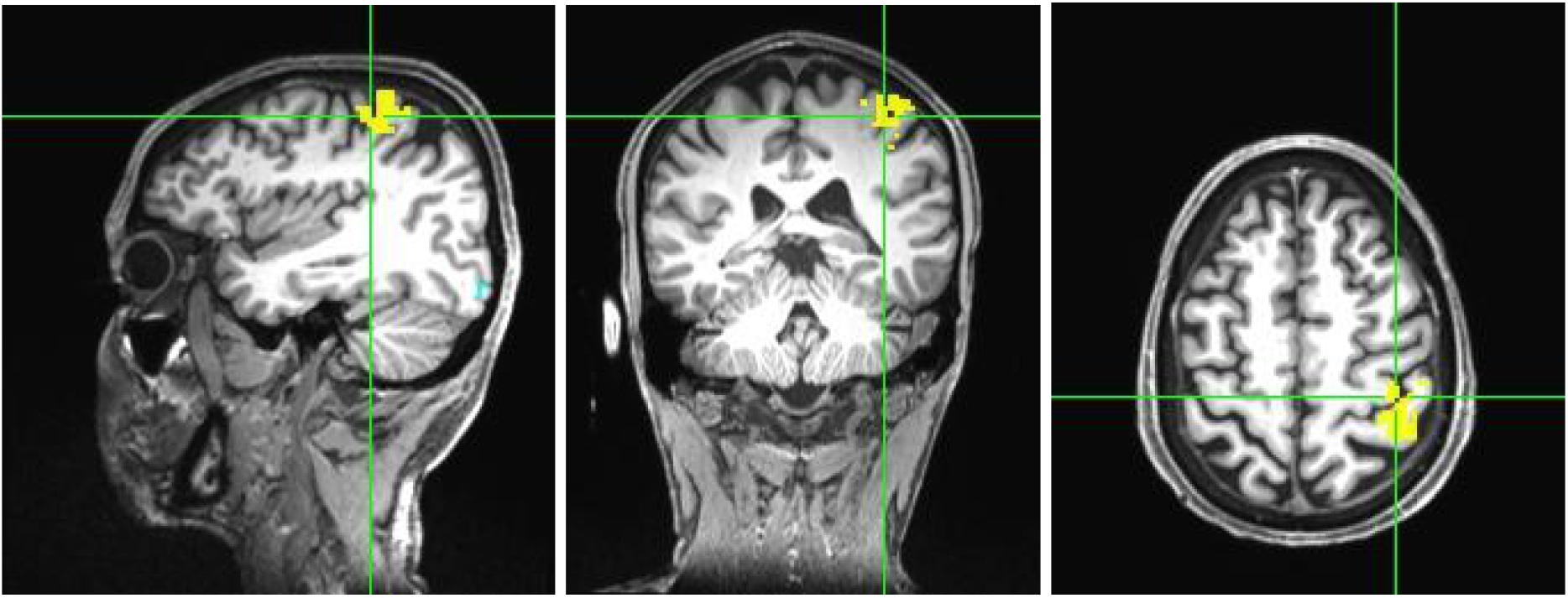
Statistical map of BOLD activity during simple sequential finger tapping with left hand. A target “hot spot” was manually selected on the cortical surface of the “hand knob” area of M1 in the precentral gyrus.

### Electrode Placement

For each participant in the imaging cohort (N=10), we first generated a current flow model using ROAST (Realistic, vOlumetric Approach to Simulate Transcranial electric stimulation)^31–33^. The model of the head was automatically generated from the T1-weighted anatomical MRI for each subject. The model was then used to determine the electrode placement that maximizes electric field at the target location in the desired orientation (example for one subject in Fig. 2a-b). To determine desired field orientation, we visually inspected the anatomical image and determined the normal vector to the cortical surface at the target in the axial scan. Polarity determined whether the vector pointed into or out of the cortical surface. For reproducibility of results without the need for costly individual MRIs, we created a single montage that provided reasonable results for all 10 participants in the imaging cohort. To this end we first selected a common desired field orientation as the mean orientation across all 10 subjects (33 degrees from to the anterior/posterior direction and 11.3 degrees from the horizontal/axial plane). We then identified for each participant the electrode montage that maximizes field magnitude for this desired orientation at the individual target. We limited total current to 4mA with a maximum of 1mA per electrode. This will always give 4 anodes and 4 cathodes^34^ but usually at different locations for each subject. Candidate locations were on the 10-10 international system. To decide on a single montage we tallied the “vote” how often a given electrode was selected (Fig. 2c). The four locations with the most “votes’’ were selected for each polarity to yield the final montage (Fig. 2d), with cathodes on F4, F2, AF4, and Fz, and anodes on P4, CP4, CP2, and P2. A reference electrode was placed at CP3 (drawing no current).

**Figure 2.**
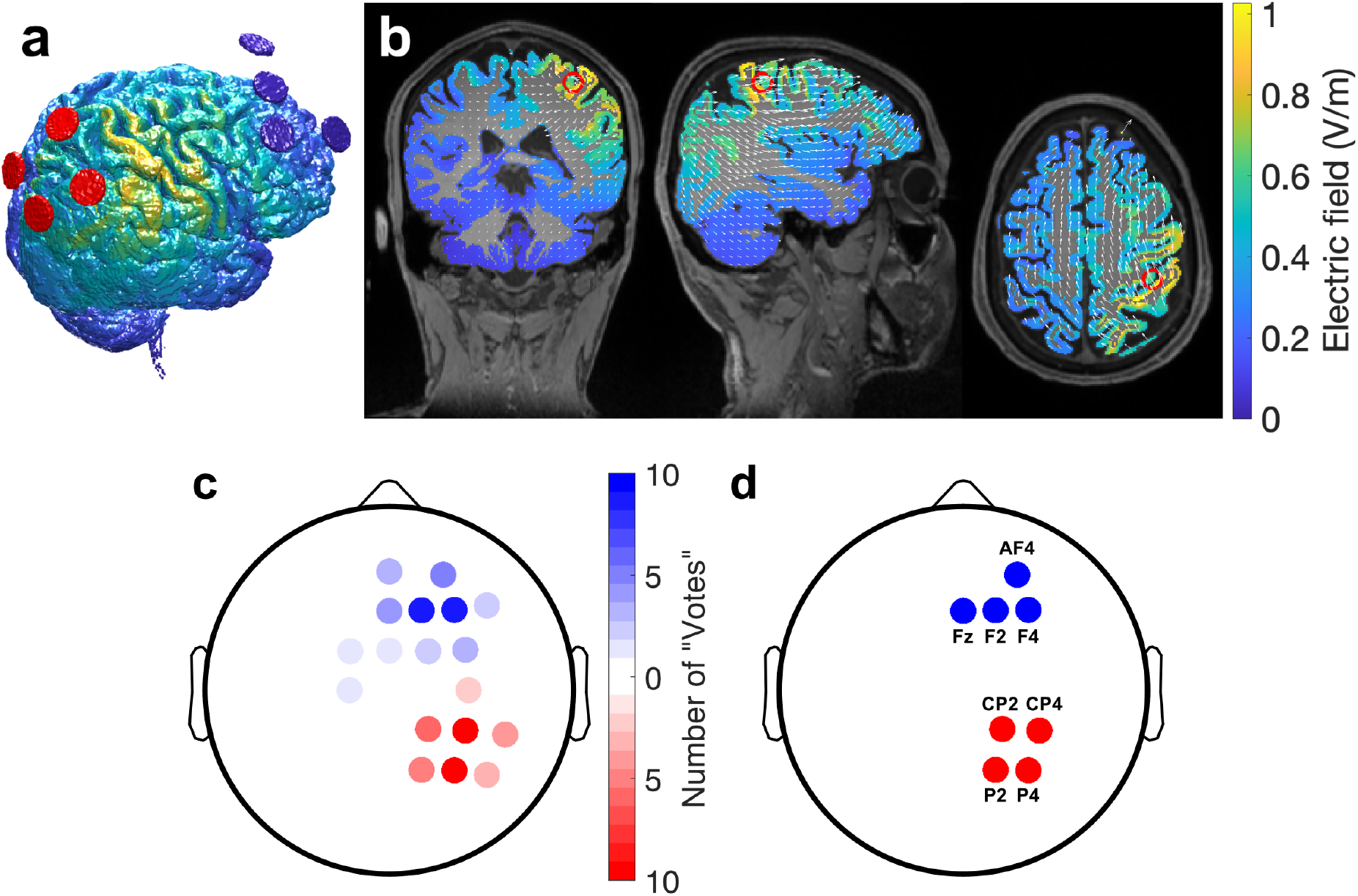
Selection of electrode montage. (a) Electric field estimated for an individual subject with a 4+4 electrode montage (red and blue circles represent 4 anodes and 4 cathode, respectively). This montage was optimized to achieve maximum intensity at the voxel in the “hand knob” with the highest fMRI activation, with current flowing from posterior to anterior direction normal to the cortical surface. (b) 2D view of estimated electric field on gray matter resulting from stimulation for the same subject in (a). White arrows represent electric field vectors and red rings mark the manually selected “hand knob” target. (c) Tally of how often an electrode location (on the 10-10 system) was selected by the optimization routine in ROAST for the 10 individual subjects models and targets (red: anodes, blue: cathodes) (d) Locations selected (with the highest tally in panel c) for a common montage that was used for all subjects during the motor sequence learning task. Anodes (red) each inject 1mA and cathodes (blue) each draw 1mA for a total of 4mA of constant current stimulation.

### Stimulation and Behavioral Task

#### Participants

For the motor sequence learning task 108 healthy, right-handed adults (52 female, 56 male, age range = 18–55 years, mean±SD = 24.6±6.42 years) provided written consent to participate in this study, under approval of the City University of New York IRB. Exclusion criteria for potential participants included any history of neurological or psychiatric disorders, traumatic brain injury, disabilities in the upper extremities, or severe visual impairment. Musicians were also excluded from the study, including professionals and amateurs who were regularly practicing a musical instrument that involves fine sequential finger movements, such as piano, guitar, violin, and trumpet, and those who had recent or extensive prior experience with such instruments but have since stopped playing. Most but not all subjects were naive to tDCS and electrical stimulation. All subjects completed the full session of experimental procedures, even though they were allowed to withdraw due to discomfort at any point during the procedure.

#### Experimental Design

In a single-blind design, participants were randomly assigned into anodal or cathodal stimulation groups (n=36 per group). Anodal stimulation serves as the active, excitatory condition. Cathodal stimulation is used as an active sham because at 4mA the sensation is noticeable and can not be reasonably shammed. To establish a baseline level of learning we subsequently tested a control group where all subjects obtained no stimulation (n=36). However, in order to simulate the stimulation environment as closely as possible, participants in the no-stimulation control group underwent the same procedures as the other two groups, including wearing a cap and having gel applied to the scalp, and they were informed that they may be stimulated. Sample size for each group was determined a priori based on performance data collected by Bönstrup et al.^35^ in G*Power 1.3^36^. Predicting a 30% increase in performance with stimulation, the effect size Cohen’s d was determined to be approximately 0.64, and with an error probability of 5% and expected 85% power on a one-sided test, we set the group sample size at 36.

#### Procedure

Stimulation was administered using a Soterix M×N-9 HD-tES System (Soterix Medical, New York, NY) with silver/silver chloride sintered ring high-definition electrodes (Soterix Medical, New York, NY) attached to a 10-10 HD-Cap (Soterix Medical, New York, NY) at the positions determined above (Fig. 2d), with conductive gel (SignaGel, Parker Laboratories, Fairfield, NJ) applied between the scalp and electrodes.

Participants were seated in front of a computer monitor and a keyboard with 4 adjacent keys labeled “1”, “2”, “3”, and “4” from left to right. The FTT described here follows the procedures detailed by Bönstrup et al.^35^, originally conceived by Karni et al.^24^ At the beginning of each FTT section, the participant was asked to place their hand (left or right) on the labeled keys (Fig. 3a). Each iteration of the task consisted of 36 continuous trials, each 20 seconds long. During the first 10 seconds of each trial a sequence of 5 digits appeared on the monitor in a MATLAB graphical user interface. Participants were instructed to press the keys corresponding to the numbers shown on screen in the order they appear in, from left to right, “as quickly and as accurately as possible”. They were to complete the sequence as many times as possible during those 10 seconds, which were followed by a 10-second rest interval. The same sequence was displayed throughout all trials in one iteration of the FTT.

**Figure 3.**
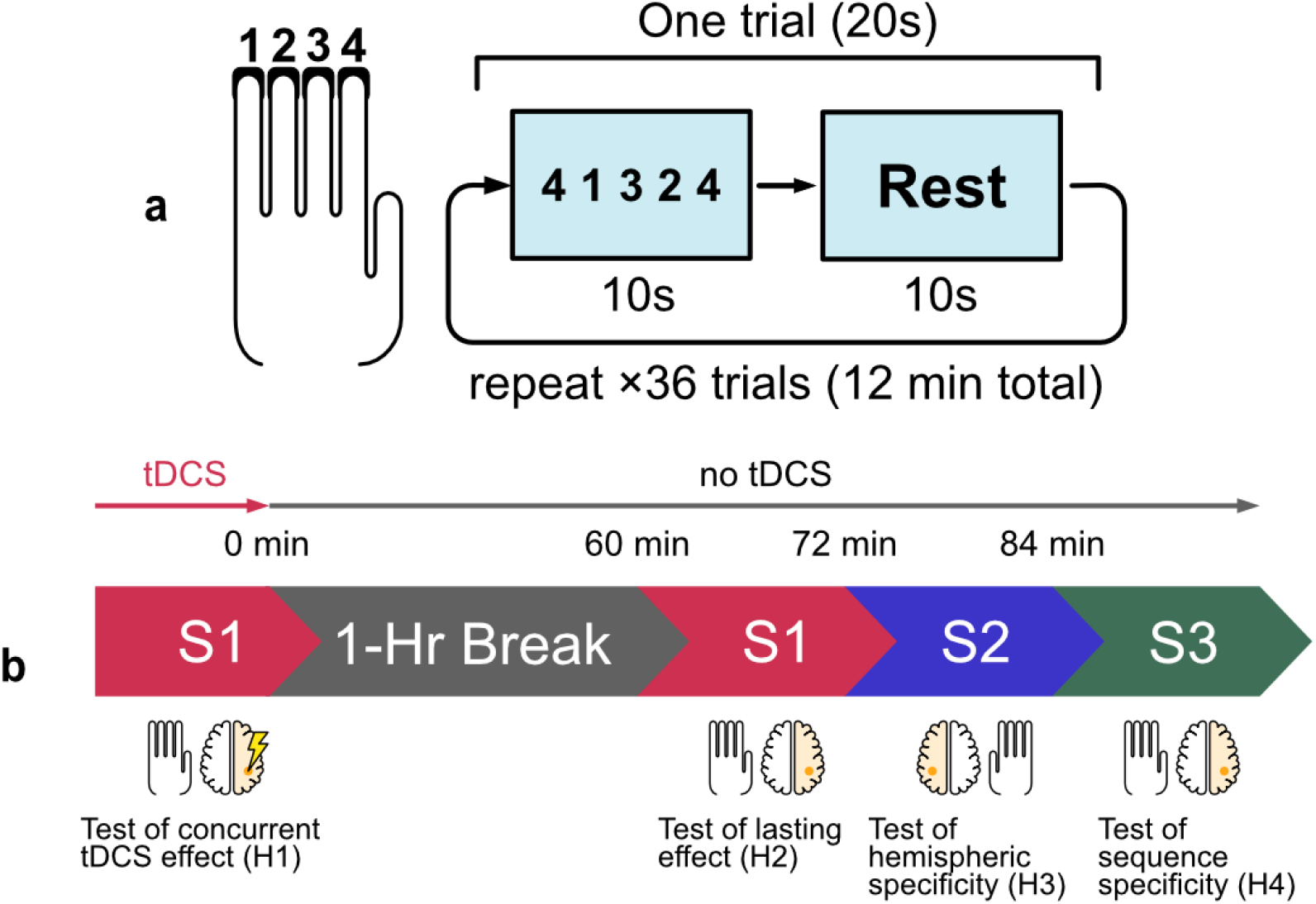
Experimental protocol. (a) Four fingers are placed on four keys labeled “1”, “2”, “3”, “4” (thumb is not used). A monitor prompts the subject to press the keys in the sequence shown. Each trial comprises a 10-second interval during which they repeatedly press the sequence as fast as possible, followed by 10 seconds of rest. Each section contains 36 trials lasting 12 minutes total. (b) An initial learning section using the left (non-dominant) hand is paired with 12 minutes of stimulation of the contralateral hemisphere (right). This is followed by a 1-hour break. The task is repeated to test for carryover effects without stimulation. Different sequences S1, S2, and S3 are used throughout different sections. S1: 4-1-3-2-4, S2: 2-3-1-4-2, S3: 3-4-2-1-3

The experiment was divided into two sessions: the main task session to test hypothesis H1 and the follow-up session to test hypotheses H2–H4 (Fig. 3b). During the initial session, participants received stimulation simultaneously while they performed the FTT with sequence S1 (4-1-3-2-4). The stimulation intensity was ramped up over 30 seconds until it reached a maximum intensity of 4mA (1mA at each electrode), at which point the participants began the task. Like the FTT, tDCS lasted for 12 minutes, after which current ramped down over 30 seconds back to zero. A visual analog scale (Wong-Baker FACES pain scale) was administered immediately after stimulation ended. Subjects were asked to rate sensation levels from 0 to 10, 10 being the most severe, at points throughout the stimulation session: the beginning, middle, and after stimulation. Sensation quality ratings at each time point were also collected, from the options: “No sensation”, “Tingling”, “Pricking/Stinging”, “Itching”, “Burning”, “Other”.

Between the two sessions the participants had a break of one hour and were free to engage in any activity (typically they engaged with their book, smartphone, personal laptop, or went out to get lunch). The followup session began with a repeat of S1 as a test of lasting learning effect. Next, subjects learned sequence S2 (2-3-1-4-2) for 12 minutes with the same FTT, but now using the right hand. The aim here was to test if there were carry-over effects to the unstimulated hemisphere. This was then immediately followed by learning sequences S3 (3-4-2-1-3) back on the left hand again with 12 minutes of FTT. The aim was to test if there were carry-over effects to a new sequence in the stimulated hand. Note that neither of these follow-up learning tasks were paired with tDCS.

#### Statistical Analysis

The primary measure of FTT performance is the number of fully correct sequences completed during each trial. Additionally, the inverse of the average time interval between keypresses within those fully correct sequences, as well as incomplete but correct sequences at the end of each trial, was taken as the tapping speed for each trial, as described by Bönstrup et al. Comparisons between groups were done using a two-sample, two-tailed t-test on the performance averaged over the entire 12 minutes of task performance. Statistical analysis was done in MATLAB and Bayes Factor calculations were done as described by Rouder et al^37^.

## Results

### Anodal stimulation improves motor sequence learning

To test H1, the primary outcome measure was the number of correct sequences completed across all trials with concurrent stimulation. The number of correct sequences combines speed and accuracy, and is therefore less susceptible to individual variations in the speed-accuracy tradeoff. The learning trajectory over the 36 trials is shown in Figure 4a. By averaging across all trials we capture early learning gains as well as late saturation of performance. The average number of correct sequences during concurrent stimulation in the anodal group was significantly higher than that of the cathodal group (Fig. 4a; Cohen’s d = 0.705, t(70) = 2.95, p = 4.30×10^−3^, planned comparison). Tapping speed was also significantly higher in the anodal group during stimulation compared to the cathodal group (Fig. 4b; Cohen’s d = 0.605, t(70) = 2.53, p = 0.0136). Thus, the subjects in the anodal group not only completed the sequence correctly more times within the learning session, but also completed each sequence more quickly.

**Figure 4.**
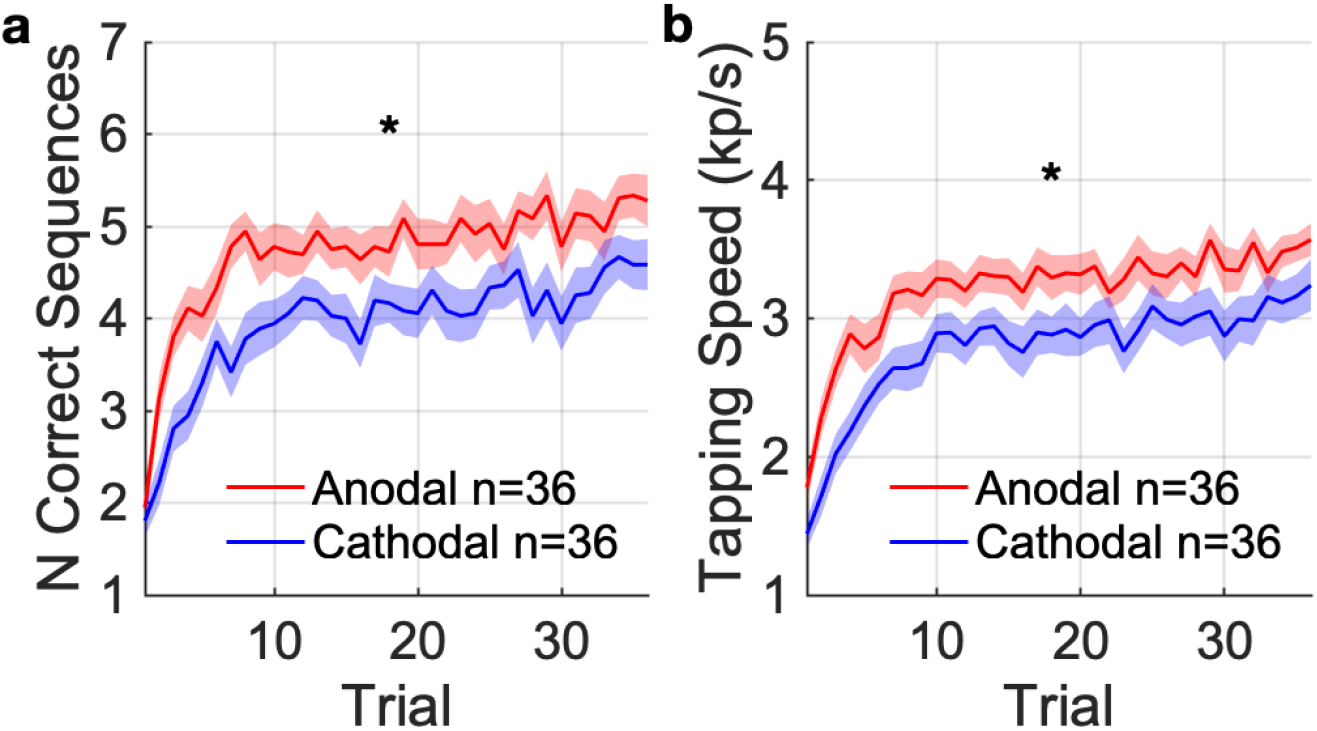
Performance in the finger tapping task with concurrent tDCS targeting contralateral motor cortex. (a) The primary outcome, measured as the number of correct sequences completed per trial (mean: solid curve, SEM: shaded area). (b) The secondary outcome, measured as the tapping speed for each trial. * indicates significant difference (p < 0.05) between anodal and cathodal groups in the average over all trials.

Prior research has shown that early performance gains do not occur during active practice of the FTT, but rather during the rest period between trials^35^. These are referred to as micro-online and micro-offline learning, respectively. The present data follow the same pattern with gains in the first 10 trials limited to micro-offline learning (see Fig. S6). Indeed, the boost in learning due to tDCS appears to be entirely constrained to boosting micro-offline learning (Table S1 for quantification and statistical tests).

### Learning gains outlasts stimulation period for at least 1 hour

The effect of polarity on performance persisted one hour after stimulation ended, in agreement with H2 (Fig. 5a). The anodal group continued to complete S1 more times than the cathodal group (t(70) = 3.13, p = 2.56×10^−3^, Fig. 5a), but the gain in speed was no longer significant (t(70) = 1.74, p = 0.0861, Fig. S1a).

**Figure 5.**
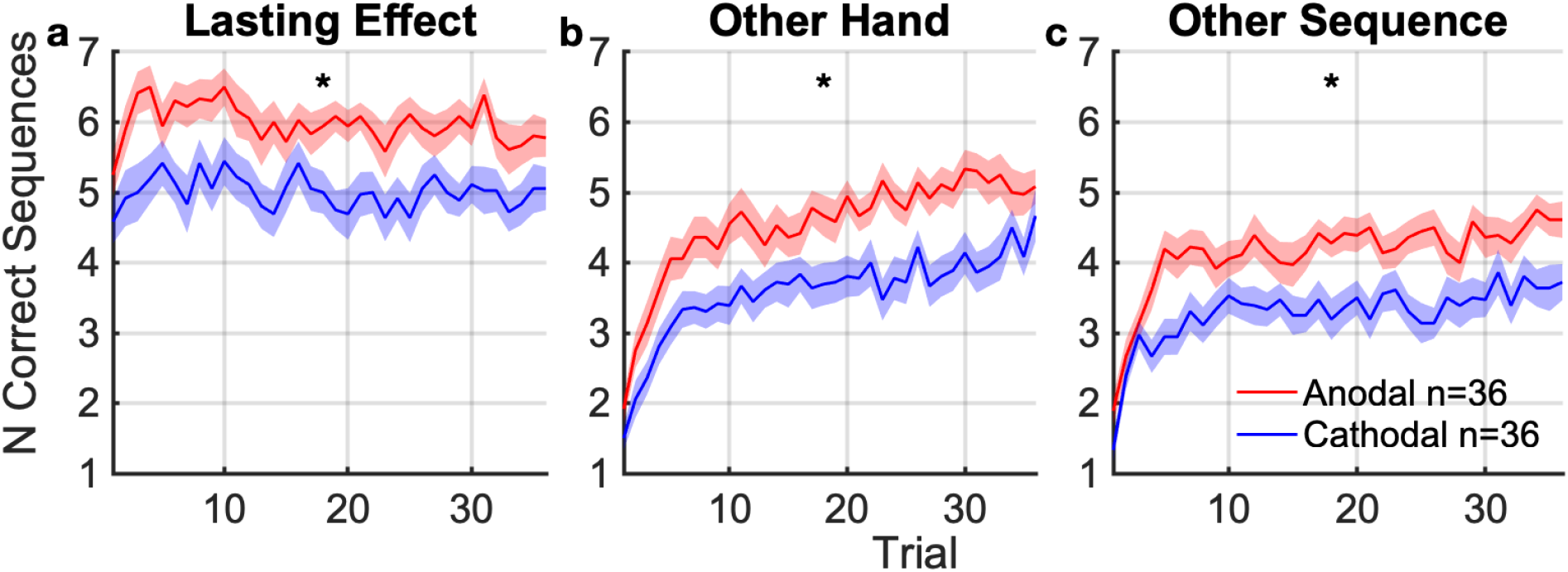
Carry-over effects of stimulation on performance and learning. Number of correct sequences completed per trial during followup tasks, averaged across subjects. (mean: solid curve, SEM: shaded area, * indicates significant difference at p < 0.05) between anodal and cathodal groups. (a) Performance with the same hand (left) and same sequence, tested 60 minutes after tDCS. This captures lasting learning effects of the targeted hemisphere and sequence. (b) Learning of a new sequence with the opposing (right) hand 72 minutes after tDCS. This captures lasting carry-over effects on learning to the other hand. (c) Learning of a new sequence with the same (left) hand 84 min after tDCS. This captures lasting carry-over effects to learning of other sequences.

### Learning gains extends to unstimulated motor sequence and unstimulated hand

Learning gains after stimulation were not specific to the stimulated hemisphere or sequence, contrary to H3 and H4. Difference between the anodal and cathodal groups carried over to learning with the right hand starting 72 min after stimulation (Fig. 5b). There was a significant difference in the number of correct sequences (t(70) = 3.28, p = 1.63×10^−3^, Fig. 5b) and a trend for tapping speed (t(70) = 1.88, p = 0.0644, Fig. S1b).

There was also a significant difference in learning of a new sequence S3 on the left hand between anodal and cathodal group 84 minutes after stimulation (Fig 5c). The gain manifested in both number of correct sequences (t(70) = 3.33, p = 1.40×10^−3^) and tapping speed (t(70) = 2.30, p = 0.0242, Fig. S1c).

Interestingly, the learning effects in the follow-up period are entirely confined to the periods of finger tapping in all stimulation conditions (Table S2). Thus the gains here should be characterized as micro-online learning - the opposite of what we observe for the initial training period, and what has been reported on this task previously ^35^.

### Anodal stimulation provided a net gain in performance over no stimulation

The follow-up experiment on a third cohort tested a no-stimulation control group (Fig. 6). The group with anodal stimulation outperformed this control group in terms of correct sequences (t(70) = 2.33, p = 0.0229, planned comparison for the follow-up experiment). Tapping speed in the anodal group during stimulation was numerically higher than the control group (Fig. S2, t(70) = 1.83, p = 0.0713). Thus, anodal stimulation boosts learning not just in contrast to cathodal stimulation but in absolute terms. Bayes factor analysis of the number of correct sequences provides moderate evidence in favor of no difference between cathodal stimulation and control (BF01 = 3.54). In general, and as expected, the no-stimulation condition falls between the cathodal and anodal condition (Fig. 6). We did not perform statistical tests for these secondary outcomes as the study was not powered to resolve these post-hoc comparisons.

**Fig 6.**
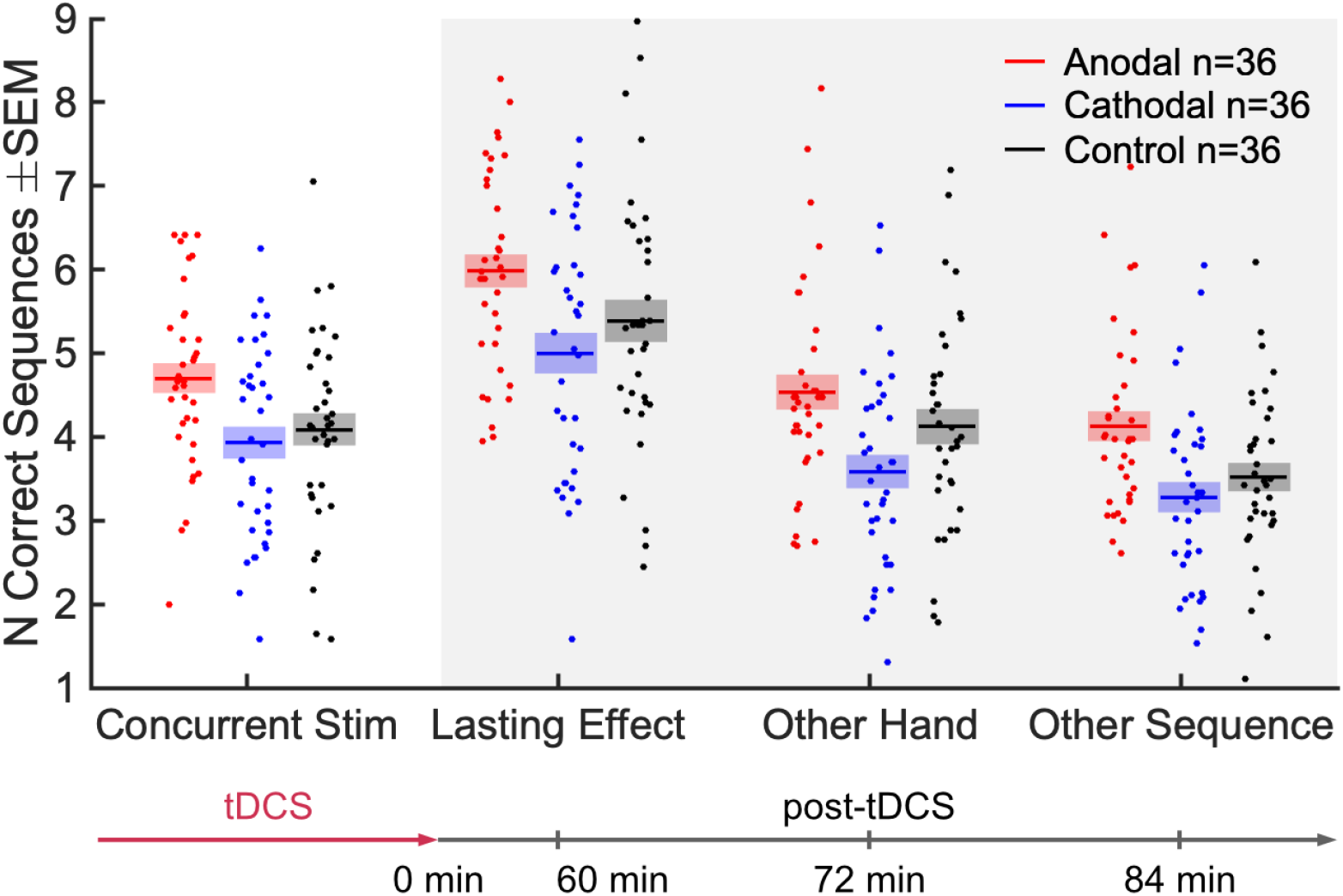
Comparison of anodal and cathodal stimulation with the follow-up control condition of no stimulation. Same data as Fig. 4a and Fig. 5, but here each point is a subject, indicating the number of correct sequences averaged over all 36 trials (mean: solid line; SEM: shaded area).

### Learning gains improved initial performance in new learning task

Thus far we have analyzed performance averaged over the entire 12 minutes of the training period, and are therefore not distinguishing gains that carry-over to initial performance vs additional gains in learning of a new sequence. We therefore test if the carry-over effects are already present at the beginning of the 12 min follow-up tests (Fig. S3). To test this we performed a two-way ANOVA on initial performance with factors of conditions (Lasting Effect, Other Hand, Other Sequence) and polarity (anodal, cathodal). We find that the initial number of correct sequences is affected by polarity (F(1) = 8.6, p = 3.7×10^−3^) but there is no interaction with task condition (F(2) = 0.15, p = 0.86). Therefore, we find a non-specific carry-over effect on initial performance. Note that initial performance in the first trial of the entire experiment differed in speed (Fig. S4) but not correct sequences (t(70) = 0.504, p = 0.616, BF01 = 4.12 in favor of null hypothesis), nor can the difference in speed predict the final outcomes (see Fig. S3-S6 for more details). This suggests that the observed polarity effects are not the results of an inhomogeneous sample of participants.

### Sensation of 4mA is tolerable and well matched with active control

The post-stimulation questionnaire records at most, moderate sensation levels at the beginning of the stimulation (Fig. 7). Sensation decreases afterwards, subsiding to very mild levels by the end of the trial. There were no significant differences in sensation ratings between the anodal and cathodal groups (BF01 = 2.59 in favor of no differences at the beginning of the trial), suggesting that the boost in learning seen in the anodal group is not the result of differing sensation. This is further supported by the lack of a performance difference between the cathodal and no-stimulation groups, despite a large difference in sensation.

**Figure 7.**
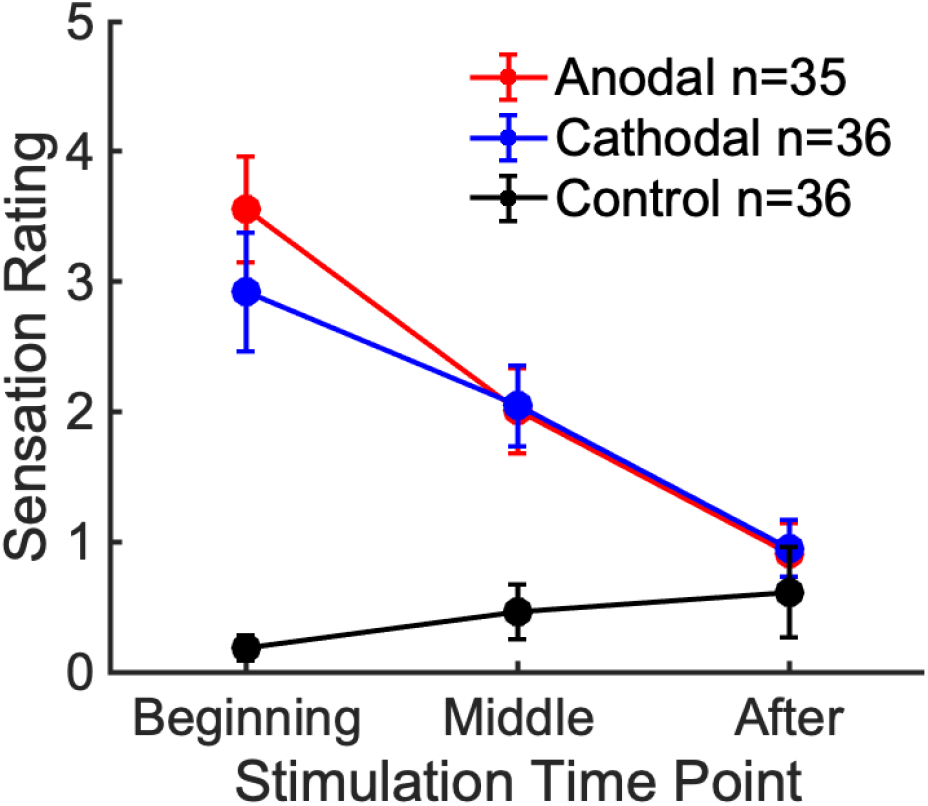
Post-stimulation VAS ratings of sensation experienced throughout different points of the stimulation session. The maximum rating participants can give is 10. A rating of ‘5’ corresponds to ‘Moderate’ sensation.

### Estimated field magnitudes and safety implications of 4+4 configuration

According to the current flow models, the proposed 4+4 electrode montage, with 4 mA total current, achieves electric fields of 0.63±0.18 V/m at the “hand knob” target on average across the 10 heads that were scanned. While this value is lower than the 0.8 V/m maximum intensity reported in previous studies with 2 mA^19^, it is important to note that those are maximum intensities across the entire brain. As seen in the coronal view (Figure 8b) E-field magnitude in gray matter is perhaps larger in the vicinity of the target under the 4+4 montage. Maximum field intensities reach up to 1.1±0.18 V/m within a 10 mm radius from the target (Fig. 8c). We compare the efficacy of this montage against other conventional tDCS montages. namely, the M1-SO (50×30×3 mm sponge anode over C4 and cathode over Fp1) and 4+1 (high-definition electrode anode over C4 and cathodes over F4, Cz, P4, and T8). The M1-SO configuration achieves 0.52±0.10 V/m on target and a maximum of 0.87±0.16 V/m within the 10mm vicinity. The 4+1 configuration archives 0.45±0.096 V/m and 0.81±0.23 V/m respectively. Aside from increasing field magnitudes, the new montage is intended to reduce current density on the scalp in order to minimize skin sensation. The highest current densities on the scalp layer are 4.1±0.17 A/m^2^, 4.6±0.27 A/m^2^, and 11±0.29 A/m^2^ for the three configurations respectively (4+4, M1-S0, and 4+1, 99.99th percentile over the entire head, Fig. 8a). Therefore, the proposed montage at 4 mA is at least an order of magnitude below the safety threshold of preclinical studies^38^.

**Figure 8.**
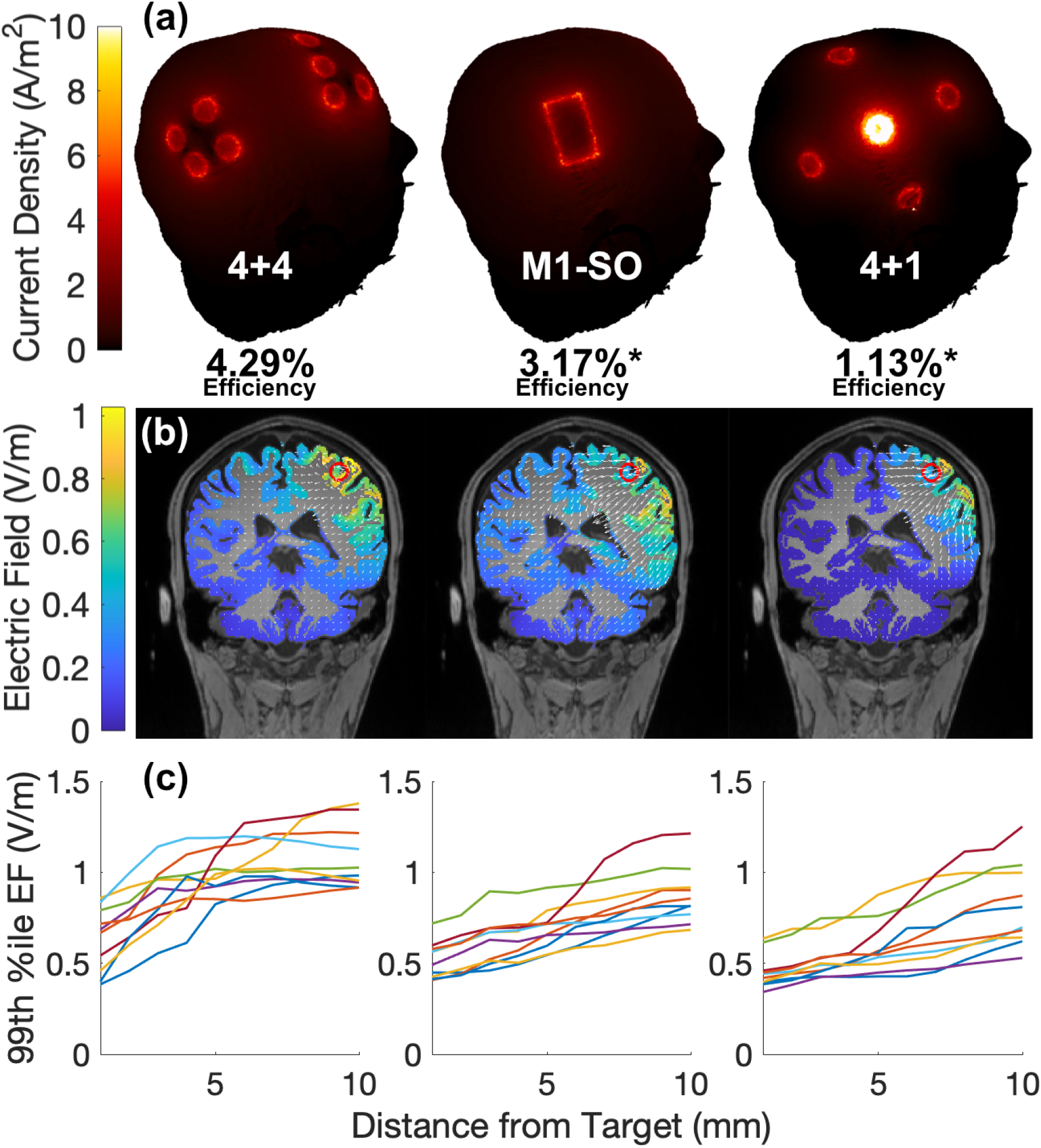
Comparisons of ROAST current flow modeling results between different tDCS configurations delivering 4mA total current. (a) 3D visualizations of current density on the skin with 4+4, M1-SO, and 4+1 tDCS montages. Stimulation efficiency value is equal to the average across 10 subjects of the ratio of current density on the target to the 99th percentile current density on the skin. * indicates a significant difference (p<0.01) from the 4+4 montage. (b) Coronal views of electric field delivered to gray matter with the respective montages in (a). The red circle marks the target and white arrows represent electric field vectors. (c) 99th percentile electric field magnitude measured in gray matter within a sphere of increasing radius around the target voxel for each subject, each represented as a separate line, with the respective montages in (a) and (b).

One way of quantifying efficiency is in terms of the tradeoff between intensity on target and discomfort, the latter of which is directly related to current density on the skin. Thus, we define stimulation efficiency as the ratio of the current density at the target over maximal current density on the scalp (Fig. 8a). We report this ratio, a unit less quantity, as a percentage. At 4.29%, the stimulation efficiency of the 4+4 montage is significantly higher than both the M1-SO (paired t-test: t(9) = 3.8, p = 3.9×10^−3^) and 4+1 montages (t(9) = 9.3, p = 6.7×10^−6^).

## Discussion

To summarize the results, anodal stimulation improves concurrent motor sequence learning with a robust effects size of 0.7. This effect outlasts the period of stimulation by at least 1 hour. The active control condition ruled out the possibility that this is the result of differing sensation levels. Importantly, the difference in performance appears to reflect a net performance gain with anodal stimulation over not stimulating at all. Sensation ratings and modeling show that distributing 4mA across 4+4 electrodes is safe, tolerable, and more effective than conventional tDCS montages.

In raising stimulation intensity to 4mA we were mindful of issues related to sensation, tolerability, and safety. Although previous efforts have demonstrated that 4mA is generally well tolerated ^17,39,40^, not all properly accounted for differences in sensation. This is especially important at 4mA, where the high intensity can no longer be ignored. Because traditional sham stimulation with ramps can provide reliably effective blinding only up to 1mA ^41–43^, we used cathodal stimulation as active control with comparable sensation. In addition, these studies delivered 4mA across a single pair of sponge or rubber electrodes. The 4+4 montage proposed here spreads out the current, passing only 1mA per electrode. This not only reduces sensation^26^, but also minimizes current density at the skin to help limit undesirable electrochemical interactions ^44^.

Computational model analysis suggests that the highest current density on the scalp under the 4+4 montage is less than half of that under an equivalent 4+1 montage. At 4mA it remains one order of magnitude lower than the threshold for tissue damage (50 A/m^2^ at 30 minutes of stimulation)^38^. Moreover, at a given current density on the skin, the 4+4 montage produces higher field intensity on target than traditional setups. Optimizing the 4+4 montage using the MRI for each individual subject can further improve this efficacy. Combined with a moderate sensation rating, 4mA tDCS through a 4+4 configuration is safe, tolerable, and effective.

The motor system has been a frequent target for tDCS due to early reports on motor cortex excitability^1^, including strong effects when controlling for various sources of individual variability^2^. However, there is a lack of standardardization and reproducibility on behavioral outcomes^10^. For example, some studies have shown an effect of anodal stimulation on FTT performance ^12,14,17^, while other studies with related motor learning tasks have found mixed effects of tDCS applied to cortex or cerebellum ^11,13,15,16,45,46^. Additionally, methodological issues complicate interpretations of these studies, such as lack of control for sensation, lack of clarity on the stimulation target, small sample sizes, and inconclusive effect sizes. Here we addressed these issues with a standard motor learning paradigm, a control for sensation, current-flow modeling, a large sample size, and conclusive effect sizes.

Selecting a stimulation target for the FTT is not always straightforward, as motor learning is a complex process that recruits the cerebellum, striatum, premotor and primary cortex (M1), supplementary motor area (SMA) and the spinal cord^7,47–54^. We chose to target M1 because motor performance during learning correlates with changes in markers of M1 activation as measured in various neuroimaging modalities ^9,24,47,55–58^. With successive learning sessions, M1 also undergoes structural changes reflected in gray matter volume increase ^59–62^. Importantly, there is a rich literature demonstrating tDCS effects on M1 excitability ^1,2^ including studies that incorporated motor learning^46^ (albeit with inconclusive results).

Although we have found a sizable effect of anodal tDCS on motor learning, the exact underlying interaction of stimulation with learning is unclear. Numerous models have proposed that multiple functionally connected networks across the various brain regions above work in parallel during motor sequence learning and their involvement changes throughout the different stages^63^. Notably, Doyon and Ungerleider describe the involvement and interaction of cortico-striatal and cortico-cerebellar circuits^64,65^. Both systems are active in the fast stage of learning, engaging the cerebellum for coordination and the striatum for sequence memory. According to Doyon et al., over the course of motor sequence learning, reliance shifts from cortico-cerebellar to cortico-striatal, when adaptive functions become less essential^64^. At the same time, active area in M1 increases later on and is positively correlated with performance^56^, so M1 appears to play a role in task representation. Due to the apparent changes in recruitment of neurons in M1, it is possible that stimulation of M1 facilitates those connections or facilitates cortico-cerebellar and cortico-striatal interaction.

The distinction between explicit and implicit sequence learning further indicates that M1 and the cortico-striatal system are especially important in the FTT used here. The FTT is considered an explicit learning task which involves not only an automation of the sequence of finger movements, but at first also requires the participant to learn the mapping from number to digits (visuomotor mapping). As opposed to an implicit learning task such as the serial reaction time task (SRTT), there is no involvement of working memory in the FTT. Comparing fMRI during both types of learning, Aizenstein et al. found that in explicit learning, M1 activity is higher during sequence learning than random key pressing, whereas with implicit learning, M1 activity is lower during sequence learning than random key pressing^66^. They also found more visual cortex, prefrontal cortex, and striatum involvement with explicit learning. It is therefore possible that having to consciously and explicitly “think” about a sequence requires M1, where it executes the logic of the movements while the cerebellum refines the movements. In addition, the parallel activation of the striatum indicates that the cortico-striatal system is more active during explicit learning than during implicit learning.

tDCS effects during the initial task were largest on micro-offline gains, during rest intervals interspersed with training bouts. This micro-offline learning has been attributed to consolidation of learning^35^. Buch et al. recently showed that it is during the rest intervals that the human central nervous system “replays” the previously practiced task^67^. In the context of this task the rate of hippocampal-neocortical replay of the practiced task predicted the magnitude of micro-offline gains. Learning in the anodal group had more pronounced micro-offline gains, suggesting an effect of anodal tDCS on the consolidation associated with this rapid replay. It would be interesting to evaluate in the future the relationship between tDCS effects and neural replay.

We note that the gain in micro-offline learning for the anodal group was offset by negative micro-online changes. We speculate that faster initial tapping speed during each trial may have caused faster fatigue in the fingers. Finally, unlike the initial task, in all three groups, learning in the follow-up tasks with S2 and S3 was dominated by micro-online learning. This unexpected shift in micro-scale learning indicates that subsequent training in the FTT may have different underlying mechanisms. This may be explained by Doyon and colleagues’ theory of motor learning, which distinguishes between an early consolidation phase and a later automatization phase, with differing underlying network mechanisms ^64^. It is possible that the change we observe after one hour of rest causes a shift between these early and later phases of learning. Regardless, this observation of a shift in learning behavior merits further investigation, irrespective of the effects of tDCS.

Based on our previous findings with DCS, this work further expands on motor learning research by examining the specificity of tDCS effects. In terms of tDCS, we observe a carryover effect across the rest period. Specifically, learning itself is boosted for both a new sequence and on the opposite hand. This appears to contradict the *in vitro* DCS work in our lab on synaptic plasticity, which suggests that stimulation has to be concurrent with training and that effects are specific to the learned task (H4)^4^. Here instead we observe that learning gains extend in time to a new motor sequence practiced without concurrent stimulation, and note that performance was improved even at baseline for the new sequence. It is possible that this unspecific boost applies to a more general skill such as the visuomotor mapping portion of the learning task, in which case we would expect a carryover to an improved baseline performance on the new sequence of the same hand. In the past, neural markers of plastic changes after motor learning were observed only for the trained hand^68,69^, suggesting hemispheric specificity (H3). Consistent with this interpretation, we only see a followup gain in baseline performance on the new sequence for the stimulated hand and not for the unstimulated hand. Conversely, there is nonetheless an overall improvement across groups in learning of new sequences over 1 hour after stimulation for both hands, even though we aimed to stimulate only one hemisphere. We draw two potential conclusions from this.

First, the stimulation may have caused a form of meta-learning, i.e. learning in general is improved after stimulation (at least up to 84 minutes, when we last tested learning). Second, this meta-learning effect is not specific to the stimulated hemisphere and not due to enhancement of visuomotor mapping, which we would have expected to be specific to the trained hand (as the mapping is different for the other hand). Another factor that may have led to unspecific gains is a lasting enhancement of attention. Future experiments would be needed and specifically designed to distinguish between these possibilities.

There could also be methodological explanations for the observed non-specific effect of tDCS on motor learning. For instance, despite randomization we could have had an inhomogeneous sample, i.e. the cohort of participants in the anodal group just happened to be better learners. Indeed, motor learning performance can be quite variable across subjects, as a result of long term training (e.g. in musicians) with concomitant increase in gray matter volume or density ^59,70^. However, a quantitative analysis of the starting performance made this possibility unlikely. Alternatively, in a single blinded experiment, the experimenter may have introduced a non-specific bias on the participants. However, given that the instructions to the participants followed a strict script, this also seems unlikely. A replication of this study on a new cohort of participants with double blinding could help address these concerns.

Nevertheless, it is safe to conclude that the effects observed were not specific to training that was concurrent with stimulation pointing to a more general neuromodulatory mechanism. For instance, it has often been argued that increased neuronal activity leads to an increase in BDNF synthesis and release, which could cause a lasting gain to plasticity, even during subsequent induction ^20,21,71^. In contrast, the mediator of DCS effects we have previously hypothesized ^4,5^ is the acute modulation in membrane potential, which can only act on synaptic plasticity during concurrent stimulation. While in-vitro experimentation is needed to address these cellular and molecular mechanisms, ultimately we require a reliable go-to experimental protocol to test behavioral learning effects in humans. Without reliable replication there is no science. It is our hope that the protocol presented here will be independently replicated by other laboratories and thus provide a firm stepping stone on the path of progress for the science of transcranial electric stimulation.

## Acknowledgements

We would like to thank Yu (Andy) Huang for his extensive support with using ROAST and his current-flow modeling expertise. We would also like to acknowledge the Magnetic Resonance Imaging Facility of CUNY Advanced Science Research Center for instrument use and technical assistance. This work was supported by the NIH through grants R21NS115018, R01DC018589 and R01NS095123.

## Conflicts of Interest

LP is listed as inventor in patents owned by CCNY, and has shares in Soterix Medical Inc.

## References

1. Nitsche MA, Paulus W. Excitability changes induced in the human motor cortex by weak transcranial direct current stimulation. J Physiol. 2000;527(3):633–639. doi:10.1111/j.1469-7793.2000.t01-1-00633.x

2. Ahn S, Fröhlich F. Pinging the brain with transcranial magnetic stimulation reveals cortical reactivity in time and space. Brain Stimul Basic Transl Clin Res Neuromodulation. 2021;14(2):304–315. doi:10.1016/j.brs.2021.01.018

3. Stagg CJ, Nitsche MA. Physiological Basis of Transcranial Direct Current Stimulation. The Neuroscientist. 2011;17(1):37–53. doi:10.1177/1073858410386614

4. Kronberg G, Rahman A, Sharma M, Bikson M, Parra LC. Direct current stimulation boosts hebbian plasticity in vitro. Brain Stimulat. 2020;13(2):287–301. doi:10.1016/j.brs.2019.10.014

5. Kronberg G, Bridi M, Abel T, Bikson M, Parra LC. Direct Current Stimulation Modulates LTP and LTD: Activity Dependence and Dendritic Effects. Brain Stimulat. 2017;10(1):51–58. doi:10.1016/j.brs.2016.10.001

6. Farahani F, Kronberg G, FallahRad M, Oviedo HV, Parra LC. Effects of direct current stimulation on synaptic plasticity in a single neuron. Brain Stimul Basic Transl Clin Res Neuromodulation. 2021;14(3):588–597. doi:10.1016/j.brs.2021.03.001

7. Dayan E, Cohen LG. Neuroplasticity Subserving Motor Skill Learning. Neuron. 2011;72(3):443–454. doi:10.1016/j.neuron.2011.10.008

8. Rioult-Pedotti MS, Friedman D, Hess G, Donoghue JP. Strengthening of horizontal cortical connections following skill learning. Nat Neurosci. 1998;1(3):230–234. doi:10.1038/678

9. Ziemann U, Iliać TV, Pauli C, Meintzschel F, Ruge D. Learning Modifies Subsequent Induction of Long-Term Potentiation-Like and Long-Term Depression-Like Plasticity in Human Motor Cortex. J Neurosci. 2004;24(7):1666–1672. doi:10.1523/JNEUROSCI.5016-03.2004

10. Buch ER, Santarnecchi E, Antal A, et al. Effects of tDCS on motor learning and memory formation: A consensus and critical position paper. Clin Neurophysiol. 2017;128(4):589–603. doi:10.1016/j.clinph.2017.01.004

11. Stagg CJ, Jayaram G, Pastor D, Kincses ZT, Matthews PM, Johansen-Berg H. Polarity and timing-dependent effects of transcranial direct current stimulation in explicit motor learning. Neuropsychologia. 2011;49(5):800–804. doi:10.1016/j.neuropsychologia.2011.02.009

12. Saucedo Marquez CM, Zhang X, Swinnen SP, Meesen R, Wenderoth N. Task-Specific Effect of Transcranial Direct Current Stimulation on Motor Learning. Front Hum Neurosci. 2013;7. doi:10.3389/fnhum.2013.00333

13. Liebrand M, Karabanov A, Antonenko D, et al. Beneficial effects of cerebellar tDCS on motor learning are associated with altered putamen-cerebellar connectivity: A simultaneous tDCS-fMRI study. NeuroImage. 2020;223:117363. doi:10.1016/j.neuroimage.2020.117363

14. Saimpont A, Mercier C, Malouin F, et al. Anodal transcranial direct current stimulation enhances the effects of motor imagery training in a finger tapping task. Eur J Neurosci. 2016;43(1):113–119. doi:https://doi.org/10.1111/ejn.13122

15. Küper M, Mallick JS, Ernst T, et al. Cerebellar transcranial direct current stimulation modulates the fMRI signal in the cerebellar nuclei in a simple motor task. Brain Stimulat. 2019;12(5):1169–1176. doi:10.1016/j.brs.2019.04.002

16. Nguemeni C, Stiehl A, Hiew S, Zeller D. No Impact of Cerebellar Anodal Transcranial Direct Current Stimulation at Three Different Timings on Motor Learning in a Sequential Finger-Tapping Task. Front Hum Neurosci. 2021;15. doi:10.3389/fnhum.2021.631517

17. Shinde AB, Lerud KD, Munsch F, Alsop DC, Schlaug G. Effects of tDCS dose and electrode montage on regional cerebral blood flow and motor behavior. NeuroImage. 2021;237:118144. doi:10.1016/j.neuroimage.2021.118144

18. Hsu G, Farahani F, Parra LC. Cutaneous sensation of electrical stimulation waveforms. Brain Stimulat. 2021;14(3):693–702. doi:10.1016/j.brs.2021.04.008

19. Huang Y, Liu A, Lafon B, et al. Measurements and models of electric fields in the in vivo human brain during transcranial electric stimulation. Brain Stimul Basic Transl Clin Res Neuromodulation. 2017;10(4):e25–e26. doi:10.1016/j.brs.2017.04.022

20. Fritsch B, Reis J, Martinowich K, et al. Direct Current Stimulation Promotes BDNF-Dependent Synaptic Plasticity: Potential Implications for Motor Learning. Neuron. 2010;66(2):198–204. doi:10.1016/j.neuron.2010.03.035

21. Ranieri F, Podda MV, Riccardi E, et al. Modulation of LTP at rat hippocampal CA3-CA1 synapses by direct current stimulation. J Neurophysiol. 2012;107(7):1868–1880. doi:10.1152/jn.00319.2011

22. Sharma M, Farahani F, Bikson M, Parra LC. Weak DCS causes a relatively strong cumulative boost of synaptic plasticity with spaced learning. Brain Stimul Basic Transl Clin Res Neuromodulation. 2022;15(1):57–62. doi:10.1016/j.brs.2021.10.552

23. Sun Y, Lipton JO, Boyle LM, et al. Direct current stimulation induces mGluR5-dependent neocortical plasticity. Ann Neurol. 2016;80(2):233–246. doi:10.1002/ana.24708

24. Karni A, Meyer G, Jezzard P, Adams MM, Turner R, Ungerleider LG. Functional MRI evidence for adult motor cortex plasticity during motor skill learning. Nature. 1995;377(6545):155–158. doi:10.1038/377155a0

25. Datta A, Bansal V, Diaz J, Patel J, Reato D, Bikson M. Gyri-precise head model of transcranial direct current stimulation: Improved spatial focality using a ring electrode versus conventional rectangular pad. Brain Stimul Basic Transl Clin Res Neuromodulation. 2009;2(4):201–207.e1. doi:10.1016/j.brs.2009.03.005

26. Borckardt JJ, Bikson M, Frohman H, et al. A Pilot Study of the Tolerability and Effects of High-Definition Transcranial Direct Current Stimulation (HD-tDCS) on Pain Perception. J Pain. 2012;13(2):112–120. doi:10.1016/j.jpain.2011.07.001

27. Reckow J, Rahman-Filipiak A, Garcia S, et al. Tolerability and blinding of 4×1 high-definition transcranial direct current stimulation (HD-tDCS) at two and three milliamps. Brain Stimulat. 2018;11(5):991–997. doi:10.1016/j.brs.2018.04.022

28. Gbadeyan O, Steinhauser M, McMahon K, Meinzer M. Safety, Tolerability, Blinding Efficacy and Behavioural Effects of a Novel MRI-Compatible, High-Definition tDCS Set-Up. Brain Stimulat. 2016;9(4):545–552. doi:10.1016/j.brs.2016.03.018

29. Desikan RS, Ségonne F, Fischl B, et al. An automated labeling system for subdividing the human cerebral cortex on MRI scans into gyral based regions of interest. NeuroImage. 2006;31(3):968–980. doi:10.1016/j.neuroimage.2006.01.021

30. Fischl B, van der Kouwe A, Destrieux C, et al. Automatically Parcellating the Human Cerebral Cortex. Cereb Cortex. 2004;14(1):11–22. doi:10.1093/cercor/bhg087

31. Huang Y, Datta A, Bikson M, Parra LC. Realistic volumetric-approach to simulate transcranial electric stimulation—ROAST—a fully automated open-source pipeline. J Neural Eng. 2019;16(5):056006. doi:10.1088/1741-2552/ab208d

32. Dmochowski JP, Datta A, Bikson M, Su Y, Parra LC. Optimized multi-electrode stimulation increases focality and intensity at target. J Neural Eng. 2011;8(4):046011. doi:10.1088/1741-2560/8/4/046011

33. Dmochowski JP, Datta A, Huang Y, et al. Targeted transcranial direct current stimulation for rehabilitation after stroke. NeuroImage. 2013;75:12–19. doi:10.1016/j.neuroimage.2013.02.049

34. Huang Y, Datta A, Parra LC. Optimization of interferential stimulation of the human brain with electrode arrays. J Neural Eng. 2020;17(3):036023. doi:10.1088/1741-2552/ab92b3

35. Bönstrup M, Iturrate I, Thompson R, Cruciani G, Censor N, Cohen LG. A Rapid Form of Offline Consolidation in Skill Learning. Curr Biol. 2019;29(8):1346–1351.e4. doi:10.1016/j.cub.2019.02.049

36. Faul F, Erdfelder E, Buchner A, Lang AG. Statistical power analyses using G*Power 3.1: Tests for correlation and regression analyses. Behav Res Methods. 2009;41(4):1149–1160. doi:10.3758/BRM.41.4.1149

37. Rouder JN, Morey RD, Speckman PL, Province JM. Default Bayes factors for ANOVA designs. J Math Psychol. 2012;56(5):356–374. doi:10.1016/j.jmp.2012.08.001

38. Bikson M, Grossman P, Thomas C, et al. Safety of Transcranial Direct Current Stimulation: Evidence Based Update 2016. Brain Stimulat. 2016;9(5):641–661. doi:10.1016/j.brs.2016.06.004

39. Khadka N, Borges H, Paneri B, et al. Adaptive current tDCS up to 4 mA. Brain Stimul Basic Transl Clin Res Neuromodulation. 2020;13(1):69–79. doi:10.1016/j.brs.2019.07.027

40. Workman CD, Fietsam AC, Rudroff T. Different Effects of 2 mA and 4 mA Transcranial Direct Current Stimulation on Muscle Activity and Torque in a Maximal Isokinetic Fatigue Task. Front Hum Neurosci. 2020;14. Accessed February 2, 2022. https://www.frontiersin.org/article/10.3389/fnhum.2020.00240

41. Gandiga PC, Hummel FC, Cohen LG. Transcranial DC stimulation (tDCS): A tool for double-blind sham-controlled clinical studies in brain stimulation. Clin Neurophysiol. 2006;117(4):845–850. doi:10.1016/j.clinph.2005.12.003

42. O’Connell NE, Cossar J, Marston L, et al. Rethinking Clinical Trials of Transcranial Direct Current Stimulation: Participant and Assessor Blinding Is Inadequate at Intensities of 2mA. PLOS ONE. 2012;7(10):e47514. doi:10.1371/journal.pone.0047514

43. Workman CD, Fietsam AC, Kamholz J, Rudroff T. Women report more severe sensations from 2 mA and 4 mA transcranial direct current stimulation than men. Eur J Neurosci. 2021;53(8):2696–2702. doi:10.1111/ejn.15070

44. Merrill DR, Bikson M, Jefferys JGR. Electrical stimulation of excitable tissue: design of efficacious and safe protocols. J Neurosci Methods. 2005;141(2):171–198. doi:10.1016/j.jneumeth.2004.10.020

45. Galea JM, Jayaram G, Ajagbe L, Celnik P. Modulation of Cerebellar Excitability by Polarity-Specific Noninvasive Direct Current Stimulation. J Neurosci. 2009;29(28):9115–9122. doi:10.1523/JNEUROSCI.2184-09.2009

46. Ambrus GG, Chaieb L, Stilling R, Rothkegel H, Antal A, Paulus W. Monitoring transcranial direct current stimulation induced changes in cortical excitability during the serial reaction time task. Neurosci Lett. 2016;616:98–104. doi:10.1016/j.neulet.2016.01.039

47. Penhune VB, Steele CJ. Parallel contributions of cerebellar, striatal and M1 mechanisms to motor sequence learning. Behav Brain Res. 2012;226(2):579–591. doi:10.1016/j.bbr.2011.09.044

48. Seitz RJ, Roland E, Bohm C, Greitz T, Stone-Elander S. Motor learning in man: a positron emission tomographic study. Neuroreport. 1990;1(1):57–60. doi:10.1097/00001756-199009000-00016

49. Doyon J, Benali H. Reorganization and plasticity in the adult brain during learning of motor skills. Curr Opin Neurobiol. 2005;15(2):161–167. doi:10.1016/j.conb.2005.03.004

50. Doyon J, Bellec P, Amsel R, et al. Contributions of the basal ganglia and functionally related brain structures to motor learning. Behav Brain Res. 2009;199(1):61–75. doi:10.1016/j.bbr.2008.11.012

51. Honda M, Deiber MP, Ibáñez V, Pascual-Leone A, Zhuang P, Hallett M. Dynamic cortical involvement in implicit and explicit motor sequence learning. A PET study. Brain. 1998;121(11):2159–2173. doi:10.1093/brain/121.11.2159

52. Hikosaka O, Nakamura K, Sakai K, Nakahara H. Central mechanisms of motor skill learning. Curr Opin Neurobiol. 2002;12(2):217–222. doi:10.1016/S0959-4388(02)00307-0

53. Vahdat S, Lungu O, Cohen-Adad J, Marchand-Pauvert V, Benali H, Doyon J. Simultaneous Brain–Cervical Cord fMRI Reveals Intrinsic Spinal Cord Plasticity during Motor Sequence Learning. PLOS Biol. 2015;13(6):e1002186. doi:10.1371/journal.pbio.1002186

54. Khatibi A, Vahdat S, Lungu O, et al. Brain-spinal cord interaction in long-term motor sequence learning in human: An fMRI study. NeuroImage. 2022;253:119111. doi:10.1016/j.neuroimage.2022.119111

55. Karni A, Meyer G, Rey-Hipolito C, et al. The acquisition of skilled motor performance: Fast and slow experience-driven changes in primary motor cortex. Proc Natl Acad Sci. 1998;95(3):861–868. doi:10.1073/pnas.95.3.861

56. Penhune VB, Doyon J. Cerebellum and M1 interaction during early learning of timed motor sequences. NeuroImage. 2005;26(3):801–812. doi:10.1016/j.neuroimage.2005.02.041

57. Kawai R, Markman T, Poddar R, et al. Motor Cortex Is Required for Learning but Not for Executing a Motor Skill. Neuron. 2015;86(3):800–812. doi:10.1016/j.neuron.2015.03.024

58. Olivo G, Lövdén M, Manzouri A, et al. Estimated Gray Matter Volume Rapidly Changes after a Short Motor Task. Cereb Cortex. Published online February 8, 2022:bhab488. doi:10.1093/cercor/bhab488

59. Sampaio-Baptista C, Scholz J, Jenkinson M, et al. Gray matter volume is associated with rate of subsequent skill learning after a long term training intervention. NeuroImage. 2014;96:158–166. doi:10.1016/j.neuroimage.2014.03.056

60. Draganski B, Gaser C, Busch V, Schuierer G, Bogdahn U, May A. Changes in grey matter induced by training. Nature. 2004;427(6972):311–312. doi:10.1038/427311a

61. Gaser C, Schlaug G. Brain Structures Differ between Musicians and Non-Musicians. J Neurosci. 2003;23(27):9240–9245.

62. Filippi M, Ceccarelli A, Pagani E, et al. Motor Learning in Healthy Humans Is Associated to Gray Matter Changes: A Tensor-Based Morphometry Study. PLOS ONE. 2010;5(4):e10198. doi:10.1371/journal.pone.0010198

63. Dahms C, Brodoehl S, Witte OW, Klingner CM. The importance of different learning stages for motor sequence learning after stroke. Hum Brain Mapp. 2020;41(1):270–286. doi:10.1002/hbm.24793

64. Doyon J, Penhune V, Ungerleider LG. Distinct contribution of the cortico-striatal and cortico-cerebellar systems to motor skill learning. Neuropsychologia. 2003;41(3):252–262. doi:10.1016/S0028-3932(02)00158-6

65. Doyon J, Ungerleider LG. Functional anatomy of motor skill learning. In: Neuropsychology of Memory, 3rd Ed. The Guilford Press; 2002:225–238.

66. Aizenstein HJ, Stenger VA, Cochran J, et al. Regional Brain Activation during Concurrent Implicit and Explicit Sequence Learning. Cereb Cortex. 2004;14(2):199–208. doi:10.1093/cercor/bhg119

67. Buch ER, Claudino L, Quentin R, Bönstrup M, Cohen LG. Consolidation of human skill linked to waking hippocampo-neocortical replay. Cell Rep. 2021;35(10):109193. doi:10.1016/j.celrep.2021.109193

68. Garry MI, Kamen G, Nordstrom MA. Hemispheric Differences in the Relationship Between Corticomotor Excitability Changes Following a Fine-Motor Task and Motor Learning. J Neurophysiol. 2004;91(4):1570–1578. doi:10.1152/jn.00595.2003

69. Floyer-Lea A, Wylezinska M, Kincses T, Matthews PM. Rapid Modulation of GABA Concentration in Human Sensorimotor Cortex During Motor Learning. J Neurophysiol. 2006;95(3):1639–1644. doi:10.1152/jn.00346.2005

70. Tomassini V, Jbabdi S, Kincses ZT, et al. Structural and functional bases for individual differences in motor learning. Hum Brain Mapp. 2011;32(3):494–508. doi:10.1002/hbm.21037

71. Podda MV, Cocco S, Mastrodonato A, et al. Anodal transcranial direct current stimulation boosts synaptic plasticity and memory in mice via epigenetic regulation of Bdnf expression. Sci Rep. 2016;6(1):22180. doi:10.1038/srep22180

