## Supplement for "Robust enhancement of motor sequence learning with 4mA transcranial electric stimulation"

**tDCS effects on tapping speed**

Tapping speed was not significantly different between anodal and cathodal conditions in the one hour followup or with the other hand (Fig. S1). However, a difference is seen in the original hand with a new sequence. Tapping speeds during all tasks were numerically higher in the anodal group than without stimulation, but the differences were not significant (Fig. S2).

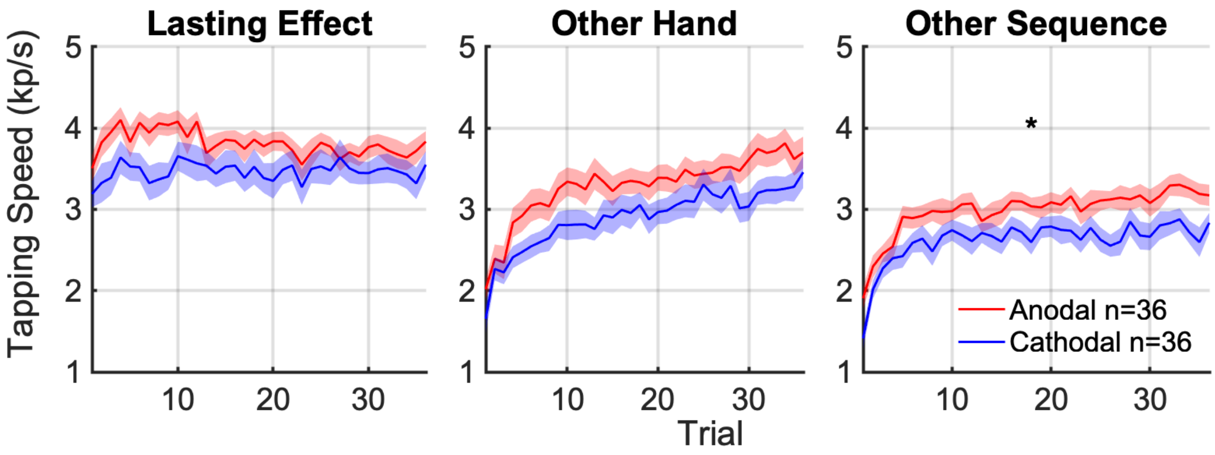


Figure S1. **Carry-over effects of stimulation on tapping speed.** Tapping speed per trial during followup tasks, averaged across subjects (mean: solid curve; SEM: shaded area; * indicates significant difference at $p < 0.05$) between anodal and cathodal groups.

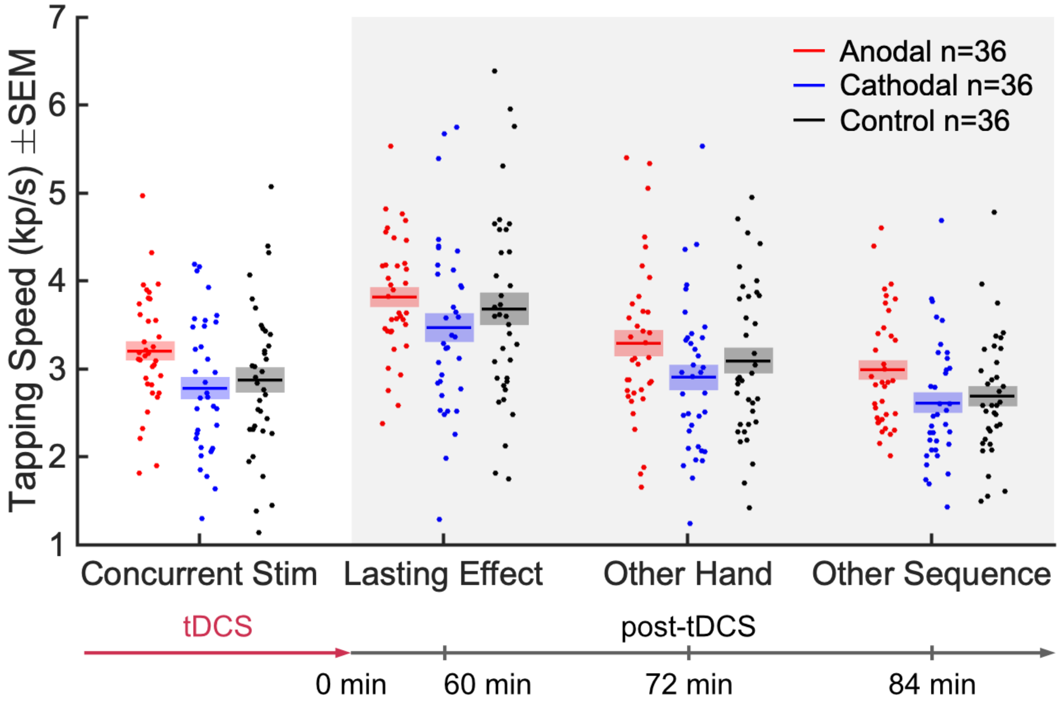


Figure S2. **Comparison of tapping speed under anodal and cathodal stimulation conditions with the follow-up control condition of no stimulation.** Tapping speed for each subject is represented as a single point indicating the tapping speed averaged over all 36 trials (mean: solid line; SEM: shaded area).

**Differences in baseline performance**

The baseline number of correct sequences during Trial 1 was not different between the anodal and cathodal groups, except in the task with a new sequence on the left hand (Fig. S3). Likewise, there is a difference in the baseline tapping speed in the final task (Fig. S4). There is also a difference in baseline tapping speed during the initial task.

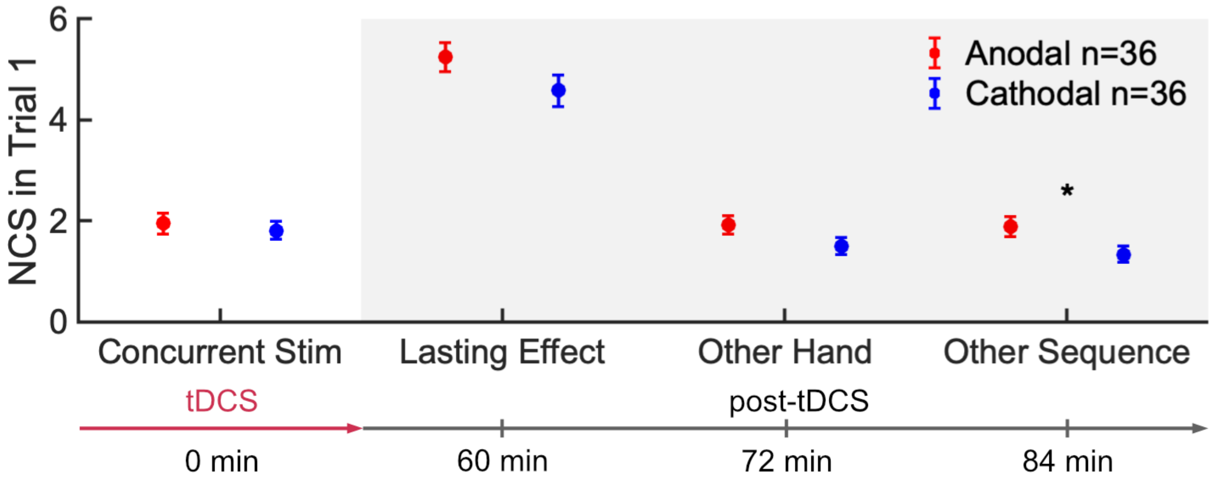

Figure S3. **Baseline number of correct sequences during the initial trial.** * indicates significant difference ($p < 0.05$) between anodal and cathodal groups. Error bars represent SEM.

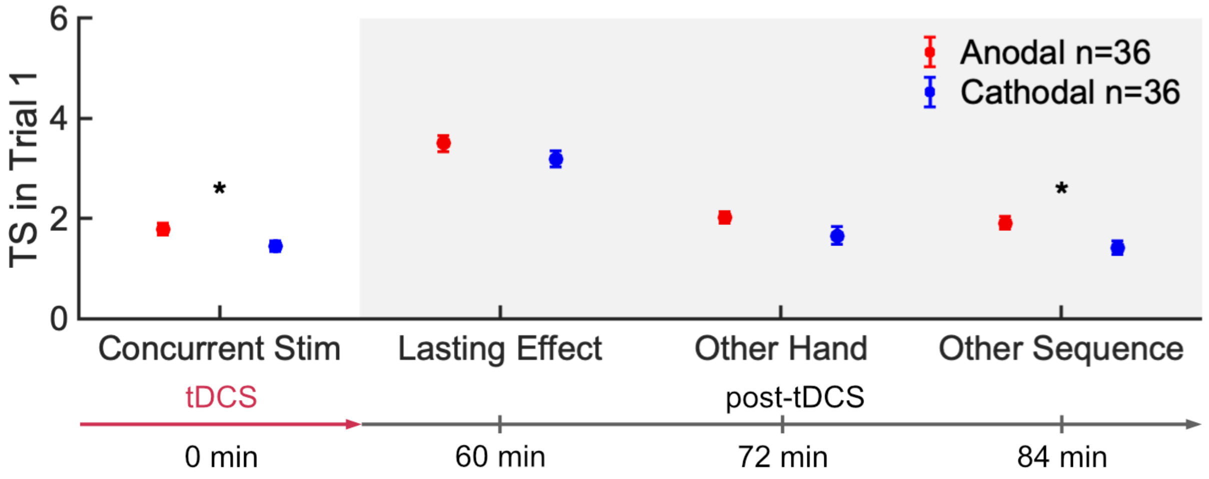


Figure S4. **Baseline tapping speed during the initial trial.** * indicates significant difference ($p < 0.05$) between anodal and cathodal groups. Error bars represent SEM.

**Dependence of final outcome on initial speed**

Is it possible that the non-specific effect of tDCS observed in the follow-up period (Fig. 5) is the result of an inhomogeneous sample? If that is the case we would expect that the anodal group performs better than the cathodal group already at the outset of the experiment. This is not true in terms of correct sequences (Fig. S3), but may have been true in terms of tapping speed (Fig. S4, $p=0.03$, uncorrected). Although these are unplanned, post-hoc and uncorrected tests, there may nonetheless be a lingering concern that we just happened to recruit a more capable group of subjects in the anodal condition. Indeed, faster tapping is positively correlated with the number of correct sequences (Fig. S5). Thus, an inhomogeneity in baseline speed may confer a benefit in the primary outcome. A simple linear regression (in the anodal and cathodal groups) between these two variables gives a slope of 0.81, and taking the initial difference in tapping speed, we predict a difference of 0.27 in the final number of correct sequences. Instead, we observed a difference of 0.69 between the two groups. Thus, the initial inhomogeneity can at most explain less than half of the actual difference observed in final performance.

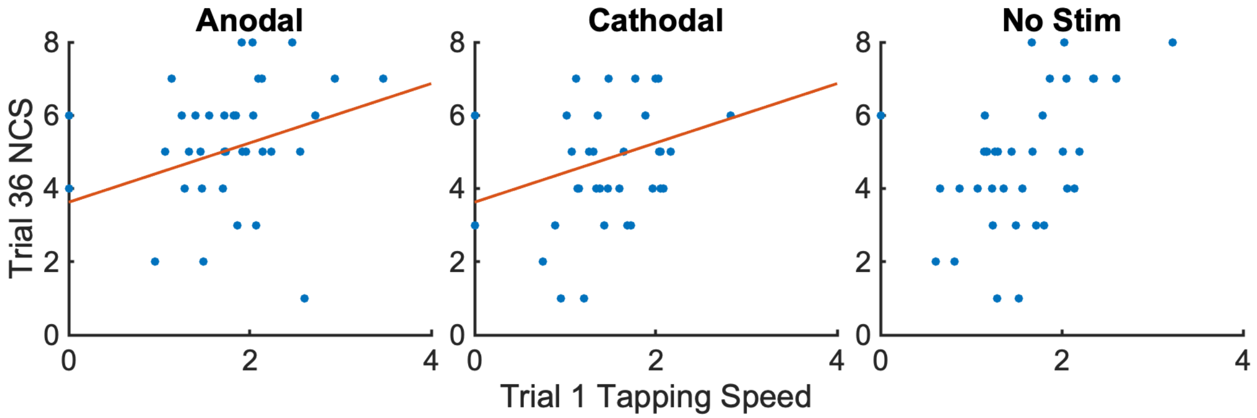


Figure S5. **Correlation of baseline tapping speed with final number of correct sequences.** Each point represents one subject. The red line shows the joint linear regression of the anodal and cathodal groups, with a slope of 0.81.

**Micro-online and Micro-offline learning**

Throughout the first 11 trials during which performance reached saturation level, micro-online changes were negligible for the cathodal group and the control group, while micro-offline changes were positive for both groups (Fig. S6 and Table S1). In contrast, we observe negative micro-online changes in the anodal group, with even more pronounced micro-offline increases indicating that the boost followed the same pattern observed in the absence of stimulation. Unlike the initial task with sequence S1, followup learning with sequences S2 and S3 was dominated by positive micro-online gains in all groups, with no micro-offline changes (Table S2).

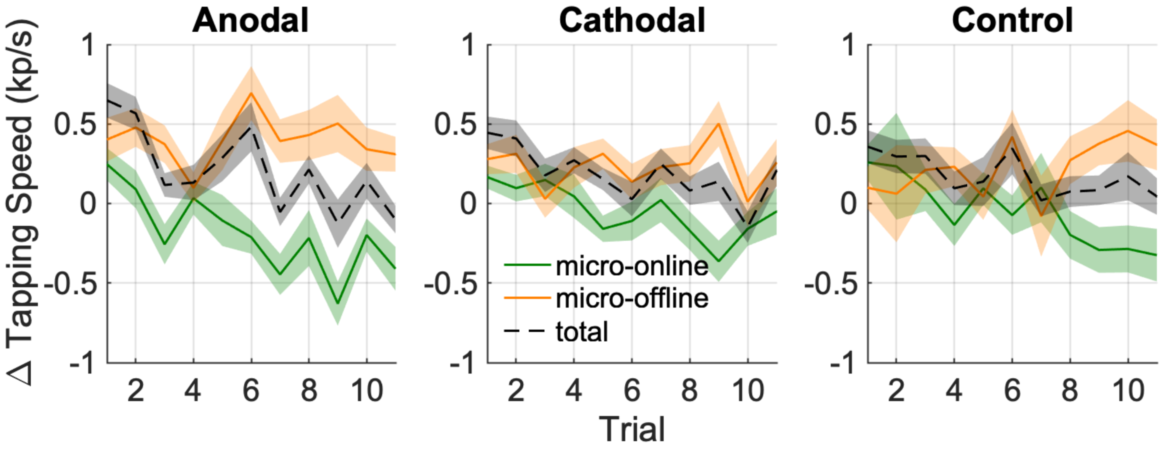


Figure S6. **Micro-online, micro-offline, and total gains in tapping speed during early learning (first 11 trials).** Micro-online gain is calculated as the change in tapping speed from the beginning of each trial to the end of the trial. Micro-offline gain is calculated as the change in tapping speed between the end of a trial and the start of the following trial. Total gain per trial represents the sum of micro-online and micro-offline gains. Shaded areas represent SEM.

Table S1. **Micro-online and micro-offline changes in tapping speed during the initial task.**

| | Micro-online learning | | | Micro-offline learning | | |
|---|---|---|---|---|---|---|
| | mean±SD | t(35) | p | mean±SD | t(35) | p |
| Anodal | -2.1±0.74 | -2.9 | $7.2\times10^{-3}$ | 4.4±0.72 | 6.1 | $5.4\times10^{-7}$ |
| Cathodal | -0.54±0.52 | -1.0 | 0.31 | 2.6±0.56 | 4.5 | $6.6\times10^{-5}$ |
| No Stim. | -0.53±0.71 | -0.75 | 0.46 | 2.5±0.76 | 3.3 | $2.5\times10^{-3}$ |

Table S2. **Micro-online and micro-offline changes in tapping speed during the followup tasks.** One-sample t-test results are shown, compared against zero. SD stands for standard deviation.

| | Micro-online learning (S2) | | | Micro-offline learning (S2) | | |
|---|---|---|---|---|---|---|
| | mean±SD | t(35) | p | mean±SD | t(35) | p |
| Anodal | 4.5±1.5 | 7.5 | $5.6\times10^{-3}$ | -2.7±1.6 | 0.42 | 0.087 |
| Cathodal | 4.3±1.4 | 7.2 | $4.6\times10^{-3}$ | -2.6±1.4 | 0.31 | 0.079 |
| No Stim. | 3.7±1.2 | 6.1 | $4.6\times10^{-3}$ | -1.9±1.2 | 0.66 | 0.14 |
| | Micro-online learning (S3) | | | Micro-offline learning (S3) | | |
| Anodal | 2.7±0.78 | 4.3 | $1.6\times10^{-3}$ | -1.1±0.82 | 0.58 | 0.19 |
| Cathodal | 2.8±1.3 | 5.5 | 0.044 | -0.97±1.3 | 1.7 | 0.47 |
| No Stim. | 2.5±0.90 | 4.3 | $9.5\times10^{-3}$ | -0.39±0.90 | 1.4 | 0.66 |
